# Reproductive proteins evolve faster than non-reproductive proteins among *Solanum* species

**DOI:** 10.1101/2020.11.30.405183

**Authors:** Leonie C. Moyle, Meng Wu, Matthew J. S. Gibson

## Abstract

Elevated rates of evolution in reproductive proteins are commonly observed in animal species, and are thought to be driven by the action of sexual selection and sexual conflict acting specifically on reproductive traits. Whether similar patterns are broadly observed in other biological groups is equivocal. Here we examine patterns of protein divergence among wild tomato species (*Solanum* section *Lycopersicon*), to understand forces shaping the evolution of reproductive genes in this diverse, rapidly evolving plant clade. By comparing rates of molecular evolution among loci expressed in reproductive and non-reproductive tissues, our aims were to test if: a) reproductive-specific loci evolve more rapidly, on average, than non-reproductive loci; ‘male’-specific loci evolve at different rates than ‘female’-specific loci; c) genes expressed exclusively in gametophytic (haploid) tissue evolve differently from genes expressed in sporophytic (diploid) tissue or in both tissue types; and d) mating system variation (a potential proxy for the expected strength of sexual selection and/or sexual conflict) affects patterns of protein evolution. We observed elevated evolutionary rates in reproductive proteins. However this pattern was most evident for female- rather than male-specific loci, both broadly and for individual loci inferred to be positively selected. These elevated rates might be facilitated by greater tissue-specificity of reproductive proteins, as faster rates were also associated with more narrow expression domains. In contrast we found little evidence that evolutionary rates are consistently different in loci experiencing haploid selection (gametophytic-exclusive loci), or in lineages with quantitatively different mating systems. Overall while reproductive protein evolution is generally elevated in this diverse plant group, some specific patterns of evolution are more complex than those reported in other (largely animal) systems, and include a more prominent role for female-specific loci among adaptively evolving genes.

## 1 Introduction

The rapid evolution of reproductive proteins has been observed across many different animal species, including in groups as diverse as marine invertebrates, insects, and mammals (Swanson and Vacquier 2002; Clark et al. 2006; Turner and Hoekstra 2008). This pattern is especially well established for proteins involved in male-specific functions, such as seminal fluid proteins, and sperm-egg and sperm-reproductive tract (RT) interactions. Although many factors can influence molecular evolutionary rates, two specific evolutionary forces—sexual selection and sexual conflict—have been proposed as primary drivers of this accelerated evolution, because both processes are expected to differentially affect reproductive traits and the proteins that underlie them. That is, traits mediating male-male competition and male-female mate choice experience unique selection to maximize mating and reproductive opportunities (Andersson 1994), and often appear to evolve rapidly between species (Ritchie 2007); therefore, the loci underpinning these reproductive traits might also be expected to display accelerated evolution and protein divergence between species (Swanson and Vacquier 2002, Clark et al. 2006, Vacquier and Swanson 2011). This inference is supported by observations of especially exaggerated protein evolution in male-exclusive and male-biased loci (collated in Dapper and Wade 2020), the sex that is usually subject to more intense sexual selection (Andersson 1994). Importantly, if sexual selection or conflict are the most critical factors driving rapid reproductive protein evolution, this pattern should be observed in other groups of organisms that also experience these selective forces. Nonetheless, whether reproductive proteins routinely evolve rapidly in non-animal species remains unclear.

Among these other groups, flowering plants (angiosperms) have numerous reproductive traits that can influence the operation and intensity of intrasexual competition and/or mate choice (Lloyd and Webb 1977, Willson 1979, Stephenson and Bertin 1983, Delph and Ashman 2006, Moore and Pannell 2011)—the two components of sexual selection. These include pollinator attraction traits, pollen competition traits, female reproductive tract (“pistil”) traits (that influence pollen performance and fertilization success), and seed maturation traits (that can be used to exercise mate choice via selective abortion) (Willson 1994, Delph and Haven 1998, Skogsmyr and Lankinen 2002). Accordingly, angiosperms may also be expected to exhibit elevated evolutionary rates in loci underpinning these traits, similar to those inferred in animals (Clark et al. 2006). Indeed, of the existing studies in angiosperms, some analyses have found evidence for faster protein evolution in reproductive loci (Szovenyi et al. 2013, Harrison et al. 2019), or greater representation of rapidly evolving genes among loci expressed in certain reproductive tissues (Gossman et al. 2016). However, others have shown more equivocal patterns (e.g. Gossman et al. 2014)—for example, sex-biased genes do not appear to evolve faster than non-sex-biased genes in several dioecious plant species (reviewed in Muyle 2019)— or have inferred greater levels of constraint on reproductive-specific proteins (e.g., Darolti et al. 2018).

These more complex findings could reflect the influence of additional factors on rates of protein evolution in plants, including the opportunity for haploid selection and the complex effects of mating system variation. In the first case, plants often express a substantial proportion of their genome during the haploid phase of the life cycle (Hafidh et al. 2016), thereby exposing these ‘gametophytic’ loci to haploid selection. In flowering plants, gametophytic traits are largely reproductive and include ‘male’ functions such as pollen tube germination, growth, and interactions with the female pistil, and ‘female’ functions such as ovule signaling (Hafidh et al. 2016, Mizuta and Higashiyama 2018). Compared to diploid-expressed loci, traits that rely on haploid gene function are expected to experience both stronger purifying selection against deleterious alleles, and elevated fixation rates (stronger positive selection) for advantageous alleles, because both deleterious and advantageous alleles will be visible to haploid selection regardless of dominance (Charlesworth and Charlesworth 1992, Walsh and Charlesworth 1992, Immler and Otto 2018, Immler 2019). Moreover, gametophytic-exclusive genes could also evolve differently from loci that are expressed in both gametophytic and sporophytic (diploid) tissue, if the latter also experience additional constraints or antagonistic effects because of their expression during both phases of the life cycle (Immler and Otto 2018).

In addition to haploid selection, plants also often exhibit substantial variation in mating system, including among closely related species. This variation is predicted to have diverse effects on the nature, timing, and strength of selection, acting both on reproductive genes specifically, and across the genome more generally. In the specific case of reproductive genes, because the mating system affects the number of reproductive partners an individual experiences, it can directly influence the intensity of intrasexual competition, the opportunity for mate choice, and the magnitude of conflict over reproductive decisions (Delph and Havens 1998; Skogsmyr and Lankinen 2002; Brandvain and Haig 2005, Clark et al. 2006, Mazer et al. 2010). Selection on traits affecting sexual competition and/or sexual conflict will be greatest in obligately outcrossing individuals, but weak or absent in obligately self-fertilizing species where the reproductive interests of both parents align perfectly (because they are the same individual). Even intermediate rates of self-fertilization can diminish the strength of sexual selection and thereby reduce, for example, the advantage of pollen competition traits (e.g. Mazer et al. 2018) and the spread of superior pollen-expressed genes (e.g. Peters and Weis 2018). Mating system—specifically variation in the frequency of selfing—is also proposed to have more global effects on patterns of selection across the genome, regardless of whether genes have reproductive functions. Because selfing reduces effective population size (*Ne*), the overall efficacy of selection—both against deleterious alleles, and in favor of advantageous alleles—is expected to be reduced in selfing compared to outcrossing lineages (Charlesworth et al. 1993, Charlesworth and Wright 2001, Mattila et al. 2018). Conversely, because selfing increases homozygosity, it can also reduce the genetic load in a population—by increasing the exposure of strongly deleterious recessive alleles to selection—as well as increase the early efficacy of selection on recessive advantageous alleles—by exposing them more rapidly to positive selection (Mattila et al. 2018). Together with specific effects on reproductive genes, these complex and sometimes antagonistic genome-wide effects might contribute to the heterogeneous inferences that have emerged from current studies of reproductive protein evolution in plants.

In this study, we examined genome-wide patterns and rates of protein evolution among four closely-related species from the diverse plant genus *Solanum*. Our aims were to compare patterns of molecular evolution in loci expressed in different classes of tissue in order to evaluate whether: a) reproductive-specific proteins evolve more rapidly, on average, than non-reproductive proteins; b) ‘male’-specific loci evolve at different rates than ‘female’-specific loci; c) genes expressed exclusively in gametophytic tissues evolve differently from genes expressed in sporophytic (diploid) tissue or in both tissue types; and d) mating system variation (a potential proxy for the expected strength of sexual selection and/or sexual conflict) affects patterns of protein evolution. In addition, to help assess underlying causes of detected patterns, we also evaluated the influence of the magnitude and breadth of gene expression on observed rates of protein evolution. Our goals were to understand the forces shaping the molecular evolution of reproductive proteins in this plant group, and whether this differs from evolutionary patterns in other protein types and/or those proposed to shape reproductive protein evolution in animal systems.

## 2 Materials and Methods

### 2.1 Study Species

Our analyses used gene expression and sequence data from each of four species within the wild tomato group (*Solanum* section *Lycopersicum*): *S. lycopersicum* (domesticated tomato; LA3475) and three wild relatives—*S. pimpinellifolium* (LA1589), *S. pennellii* (LA0716), and *S. habrochaites* (LA1777) (Figure S1). (LA#### refers to the specific germplasm accession ID, as described in tgrc.ucdavis.edu, from which all or most of the data were obtained; Table S1.) This group is a clade of 12 closely-related diploid wild species along with the domesticated tomato, within the hyperdiverse (~1300 species) plant genus *Solanum* (Sarkinen et al. 2013). All 12 wild tomato species arose within the last ~2.5 million years, consistent with recent rapid speciation in this clade (Pease et al. 2016a).

Mating system varies substantially across species in *Solanum* (Goldberg et al. 2010) and our four focal species differ in their mating system, most notably in whether species are historically self-incompatible (SI: *S. pennellii* and *S. habrochaites*) versus self-compatible (SC: *S. lycopersicum* and *S. pimpinellifolium*). These species also have large estimated differences in their genetic effective population sizes (Ne), that are consistent with these SI vs SC mating system differences. For example, prior estimates of average heterozygosity from transcriptome-wide data are 0.28% and 0.245% for *S. pennellii* and *S. habrochaites* accessions respectively, and 0.04% for *S. pimpinellifolium* accessions (Table S3 in Pease et al. 2016a). Heterozygosity is even lower in the domesticated tomato (*S. lycopersicum*) (e.g., Rick and Fobes 1975). In *Solanum*, genetic self-incompatibility (the inferred ancestral state) is based on molecular interactions between proteins expressed in growing pollen tubes and pistil-expressed SRNase and other known proteins (Cruz-Garcia et al. 2003; Takayama and Isogai 2005). SI is observed as arrest of pollen tube growth in the female reproductive tract (the pistil). Self-compatible lineages have lost function in one or more SI-mediating proteins (Stone 2002), thereby allowing individuals to be fertilized by their own pollen, in addition to outcrossing. As with other angiosperms (Sicard and Lenhard 2011), transitions to SC within *Solanum* are often followed by other morphological changes in floral size and structure, that further increase the frequency of self-pollination and reduce the number/diversity of mating partners (Rick 1979). In our four focal species, SI species have larger flowers, longer reproductive tracts, and more pollen production on average, in comparison to SC species (Vosters et al. 2014), even though the transition to SC likely occurred within the last ~0.5MY (the estimated age of the clade containing the two SC species here; Pease et al. 2016a). Both differences in the capacity to reject self pollen (i.e. SI versus SC), and these morphological differences, could have large effects on the average diversity of mating partners, and therefore the opportunity for and magnitude of intraspecific sexual selection and sexual conflict (Willson 1994, Delph and Havens 1998, Clark et al. 2006). In particular, while cross-pollination rates in SI species must be 100%, these have been estimated to range from 0-40% in wild SC species, including *S. pimpinellifolium* (e.g., Rick et al. 1977, 1978).

### 2.2 Quantifying gene expression and determining tissue-specificity

Our analyses used gene expression (RNAseq) data collected individually from leaf (up to five developmental stages), root, seed, vegetative meristem, stem, flower (several stages), style, pollen, and ovule tissues in each accession. RNAseq data were obtained from seven publically available SRA projects (Table S1), mostly drawn from three previously published analyses (Koenig et al. 2013, Ichihashi et al. 2014, Pease et al. 2016b), except data from ovule tissue which has not been previously published. Note that style and pollen data were not available specifically for *S. pimpinellifolium*. Sources of each dataset are described in detail in Table S1. Procedures for generating the ovule RNAseq data are described in the supplementary text. Each tissue had 1-6 replicates (generally 3) per accession (Table S1).

For each library within each tissue, we processed the raw reads by filtering adapter sequences and low quality bases with TRIMMOMATIC (Bolger et al. 2014). Reads were then mapped against the tomato reference genome (ITAG 2.4) using STAR (Dobin et al. 2013), with default settings excluding alignments with >10 mismatches. Numbers of reads mapped onto genic regions were estimated with FEATURECOUNTS (Liao et al. 2014). We normalized the read counts from each library by calculating TPM (transcripts per million) and then calculated the mean normalized read counts for each tissue of each species.

We first classified loci into three general classes of gene—reproductive (RP), vegetative (VG), and general (GR) (i.e., expressed in both reproductive and non-reproductive/vegetative tissue types). Reproductive genes were required to be expressed (TPM>2) in at least one reproductive tissue (i.e. style, pollen, ovule), and to have no or trace expression (TPM <0.5) in any of the remaining tissues (i.e. leaf, root, stem, seed, root, vegetative meristem), in at least two of the three investigated species for which we had all tissues available (i.e. *S. lycopersicum*, *S. pennellii*, and *S. habrochaites*). (Because *S. pimpinellifolium* lacked some of the tissue types in our RNA-seq data, we did not include this species in the filtering/determination of tissue-specificity.) Vegetative genes were defined similarly by requiring expression in at least one vegetative (non-reproductive) tissue, and no/trace expression in any reproductive tissue. Generally-expressed (GR) genes were expressed in at least one reproductive tissue and also in at least one vegetative tissue.

Using similar criteria, we also identified a set of tissue-specific genes, defined as those only expressed in one specific tissue (TPM>2) and not in any other tissue (TPM<0.5) in at least two out of the three species for which we had all tissue types (i.e. *S. lycopersicum*, *S. pennellii*, and *S. habrochaites*). For these analyses, we focused on specific tissues (i.e. leaf, root, stem, seed, vegetative meristem, style, pollen, ovule) for which we had sufficient sampling for at least three species (Table S1). Finally, using the same criteria and the expression domains in the three species for which we had all tissues represented, we also classified transcripts as gametophytic-exclusive (pollen and/or ovule expression only) versus sporophytic-exclusive versus both gametophytic and sporophytic (pollen and/or ovule expression, and at least one other diploid tissue), as well as ovule-specific versus expressed in both ovules and at least one sporophytic (diploid) tissue, and pollen-specific versus expressed in both pollen and at least one sporophytic (diploid) tissue.

### 2.3 Estimating rates of protein evolution

As with prior studies (Clark et al. 2006 and references therein), we used dN/dS—the ratio of the number of non-synonymous substitutions per non-synonymous site to the number of synonymous substitutions per synonymous site—as our estimate of protein evolution. Although dN/dS is usually <1—because most functional loci experience some constraint on non-synonymous changes—larger values of dN/dS are consistent with faster protein evolution. Note that estimates of dN/dS can be influenced by intraspecific polymorphism when it is high compared to fixed differences between species (see Hahn 2018, Chapter 7). However, estimates of heterozygosity within each of our wild accessions are small compared to divergence (e.g. dS) estimates across these species (Supplementary text), indicating that most of the SNPs contributing to our estimates of dS, dN, and dN/dS, are fixed differences.

We estimated dN/dS for each locus in our dataset, to calculate mean rates of protein evolution in different classes of genes, and to identify the set of loci with estimated dN/dS>1—a pattern consistent with positive selection for protein evolution at that locus. In addition, to compare the overall dN/dS for loci on SI versus SC branches, we compared estimates from sequences that were concatenated across groups of genes (described below). Note that low transcript coverage reduces the power to accurately call SNPs at heterozygous sites and can thereby introduce errors into inferred sequences and statistical estimates based on these; this can be a concern in low coverage RNA-seq data. However, the average per locus raw read count in our dataset ranges from ~160 to >800 (Table S2), indicating that we have ample power to accurately call SNPs with these transcript data.

To generate the sequence alignments for these analyses, we combined the RNA-seq data used for quantifying gene expression (above) with whole-transcriptome data from the same four accessions previously generated in Pease et al. (2016a), and with publicly available DNA-seq data (Table S3); the latter datasets allowed us to augment our RNAseq data with additional sequences for loci whose expression was not detected for one or more of our four species. These reads were aligned against the tomato reference genome by STAR, with defaults excluding alignments with >10 mismatches, as with the previous reference-aligned single copy ortholog dataset (Pease et al. 2016a) The aligned data were processed using SAMTOOLS followed by MVFTOOLS to generate the sequence alignments of orthologous genes. We only retained transcript sequence alignments longer than 200 bps after removing gaps and indels, leaving us with 21,216 sequence alignments (loci) for downstream analyses.

For each locus, we used PAML (Yang 2007) to estimate dN/dS across the branches of a given phylogenetic topology among the four investigated species, using the input species tree ((*lycopersicum, pimpinellifolium*), (*pennellii*, *habrochaites*)) which reflects the known species topology (Figure S1). To do so we used the one-ratio (M0) codeml model in PAML, that generates a single dN/dS estimate across all branches for each locus (that is, dN/dS is fixed at one ratio for each gene). Outputs from this model were used to compare the mean and distribution of dN/dS for genes among three broad categories (reproductive (RP), vegetative (VE), and general (GR)), as well as between each of six tissue-specific groups classified by their expression in only a single tissue (pollen, style, ovule, leaf, root, or stem; vegetative meristem and seed were excluded from the latter analyses because they had no exclusively tissue-specific genes), and between loci that had gametophytic versus sporophytic, or gametophytic+sporophytic, domains of gene expression. We also compared loci expressed in ovules only versus expressed in ovules and at least one sporophytic (diploid) tissue, or in pollen only versus expressed in pollen and at least one sporophytic tissue.

Second, in addition to locus-by-locus dN/dS comparisons, we also evaluated overall dN/dS estimates along SC branches in comparison to those along SI branches. Among our four species, the two self-compatible species (*S. lycopersicum* and *S. pimpinellifolium*) are most closely related to each other, and the lengths of SC branches (those leading to *S. lycopersicum* and *S. pimpinellifolium*) are much shorter than the length of SI branches (those leading to *S. pennellii* and *S. habrochaites*) (Figure S1). As a result, branch lengths (especially for SC lineages) were too short to reliably estimate lineage-specific dN/dS for each locus individually. Therefore, for these analyses we concatenated all loci within each class of genes to be compared, and then estimated dN/dS across the concatenated set of loci for SI and SC branches separately. For example, for reproductive (RP) loci, we concatenated all RP loci and estimated dN/dS across the entire concatenated set, allowing for different dN/dS estimates (variable rates) on SI versus SC branches. Because of the small numbers of loci concatenated for some of our individual tissues, we limited SI vs SC branch-specific estimates to our three broad classes of loci (RP, VG, and GR) only.

Because the phylogeny of our species contains one internal branch, for these analyses we used two alternative multi-rate (M2) codeml models in PAML. For the first, three-ratio, model we allowed the internal branch to have a rate that differed from the tip branches leading either to SC species, or to SI species. For the second, two-ratio, model, we classified the internal branch as ‘SI’ (i.e. dN/dS was estimated separately for tip branches leading to SC species, versus a background rate for the tip branches leading to SI species plus the internal branch). This approach was used to estimate branch-specific rates (dN/dS estimates) for SI versus SC branches for each of our three general classes of loci (RP, VG, and GR). Note we report three-ratio results in the main text, and two-ratio results in the supplement.

### 2.4 Comparing protein evolution rates

All statistical analyses were performed in R version 3.6.3 (R Core Team 2015). For all comparisons, we first removed genes showing unusually high values (dN/dS >10) from the paml output, as these were due to estimation limitations within paml (e.g. a lack of synonymous substitutions across the branch(es) used for estimation) or otherwise from poor sequence quality or alignment.

To compare the mean rate of per locus protein evolution (dN/dS) between different classes of loci, we fit generalized linear models (GLMs). Shapiro-Wilk tests of normality and quantile-quantile plots were used to assess normality in the distribution of dN/dS; in all categories, dN/dS was heavily skewed towards lower values and zero-inflated (reflecting the fact that most expressed genes are under purifying selection). To accommodate this in our models, we assume a gamma residual distribution with an identity link function, rather than the standard Gaussian assumption, after confirming the suitability of this distribution with maximum likelihood (using the fitdistrplus R package). Following GLMs, to compare the mean estimated dN/dS between specific classes of loci, we made pairwise comparisons using Tukey post-hoc tests, adjusting p-values for multiple comparisons. Specifically, post-hoc test p-values were adjusted using the default ‘single-step’ method in the R package ‘multcomp’ (Hothorn et al. 2008), which adjusts p-values based on the joint normal or t distribution of the linear function. In addition, we compared the proportion of positively selected genes (i.e. dN/dS>1) between different classes of loci by performing Chi-Squared tests of independence and, for specific tests of enrichment in single categories (e.g., in RP relative to GR), one-sided Fisher’s exact tests.

To assess evidence for differences in dN/dS between mating systems, we aimed to compare dN/dS estimates from concatenated data, as calculated separately for SI and SC branches. Both biological and technical factors make these comparisons challenging to interpret. Biologically, overall differences in Ne and therefore in the expected efficacy of positive and purifying selection, could generate global differences in rates of protein evolution between SC and SI lineages (see Introduction), that are unrelated to their potential differences in the historical strength of sexual selection. Technically, in our specific dataset, the two SC species are much more closely related than the two SI species, so there are large differences in branch lengths (number of substitutions) over which we are estimating dN/dS values for SI versus SC branches. Therefore, to assess whether exclusively reproductive (RP) loci had a different pattern of molecular evolution on SI versus SC branches in comparison to other loci, we compared the observed difference in dN/dS between SI and SC branches for RP genes to an estimate of the ‘baseline’ genome-wide difference in dN/dS between SI and SC branches. This baseline was described by the SI vs SC difference in our set of generally expressed genes (GR), which is expected to be due to factors unrelated to variation in sexual selection. Because we have only a single estimate (from the concatenated data) for each class of gene for each branch type, standard parametric tests can’t be used. Instead we used bootstrap resampling to estimate the variance around the mean SI vs SC difference in dN/dS in the GR dataset, and then to evaluate whether the observed SI vs SC difference in RP genes falls outside the range of this variance (i.e. is larger or smaller than the 95% CI of this distribution). For each bootstrap replicate (N=1000 replicates), we re-sampled 500 loci at random from the GR dataset, estimated dN/dS for SI and SC branches and the difference between these two estimates. Our observed dN/dS difference (SC-SI rate) was compared to the resulting distribution of 1000 replicate dN/dS differences from GR loci.

Finally, to confirm that detected differences in dN/dS values between classes of genes were not driven by differences in rates of synonymous mutation—which, for example, can systematically vary according to genomic location or nucleotide content—we also evaluated differences in dN and dS separately for these classes of genes (Supplementary text).

### 2.5 Assessing the effect of gene expression level on molecular evolutionary rates

Numerous studies have shown that rates of protein evolution can be influenced by the magnitude and breadth of gene expression, such that highly expressed genes and/or genes expressed in a broader number of tissues tend to have systematically lower estimated dN/dS (Meisel 2011). To assess the effect of the magnitude and breadth of gene expression on our inferences, we also compared average dN/dS values between genes expressed in different numbers of tissues (ranging from 1 (tissue-specific loci) through to 8 (expressed in all tissues examined)), using the criteria that a gene be expressed >2 TPM in a specific tissue in all species for which we had data on that tissue-type. Within each of the broad classes of RP, VG, and GR loci, we also assessed the quantitative relationship between mean gene expression level (averaged across all tissues in which a locus was expressed) and estimated dN/dS. For this analysis we constructed a multiple regression model of dN/dS (square root transformed to approximate normality), including both mean gene expression (TPM), tissue class (RP, VG, or GR), and their interaction term as independent variables.

### 2.6 Assessing enrichment of functional categories among adaptively evolving loci

To test for functional enrichment among our loci with dN/dS > 1, we performed a Gene Ontology (GO) term enrichment analysis using Panther version 14 (Mi et al., 2019; Ashburner et al., 2000; The Gene Ontology Consortium, 2019). We performed separate tests for GR, VE, and RP tissue categories, as these classes had large sample sizes appropriate for tests of enrichment. Loci were evaluated using both the biological process and molecular function annotation data sets, and significance was determined using Fisher’s Exact tests with false discovery rate correction. False discovery rates were calculated using the Benjamini-Hochberg procedure, using a critical value of 0.05 to filter results. All statistics were calculated using the automated PANTHER Overrepresentation Tests.

## 3 Results

### 3.1 Reproductive genes evolve at modestly faster rates, and have proportionally more genes under positive selection

Reproductive (RP) genes had an elevated mean rate of protein evolution compared to loci expressed in both reproductive and non-reproductive (general/GR) tissues (Posthoc Z=2.548, p = 0.0108) (Figure 1; Table 1; Table S4, Table S5). The mean dN/dS for vegetative-specific (VG) loci fell between these two other classes of genes and did not differ from either of them (Table 1). These observed differences were not driven by variation in synonymous mutation rates (dS) between these classes of genes (Supplementary text S3; Figure S2, Table S6). In addition, the proportion of genes with dN/dS>1 differed by group (Table 1). A significantly greater proportion of reproductive loci (28 of 670, or ~4.17%) are inferred to be evolving adaptively compared to general loci (2.45%) (Fisher’s exact test, p=0.0067; Table 1, Table S5). The proportion of vegetative loci evolving adaptively (2.81%) was marginally lower than reproductive loci (Fisher’s exact test, p=0.093), and did not differ from general loci (Table 1; Table 4).

**Figure 1:**
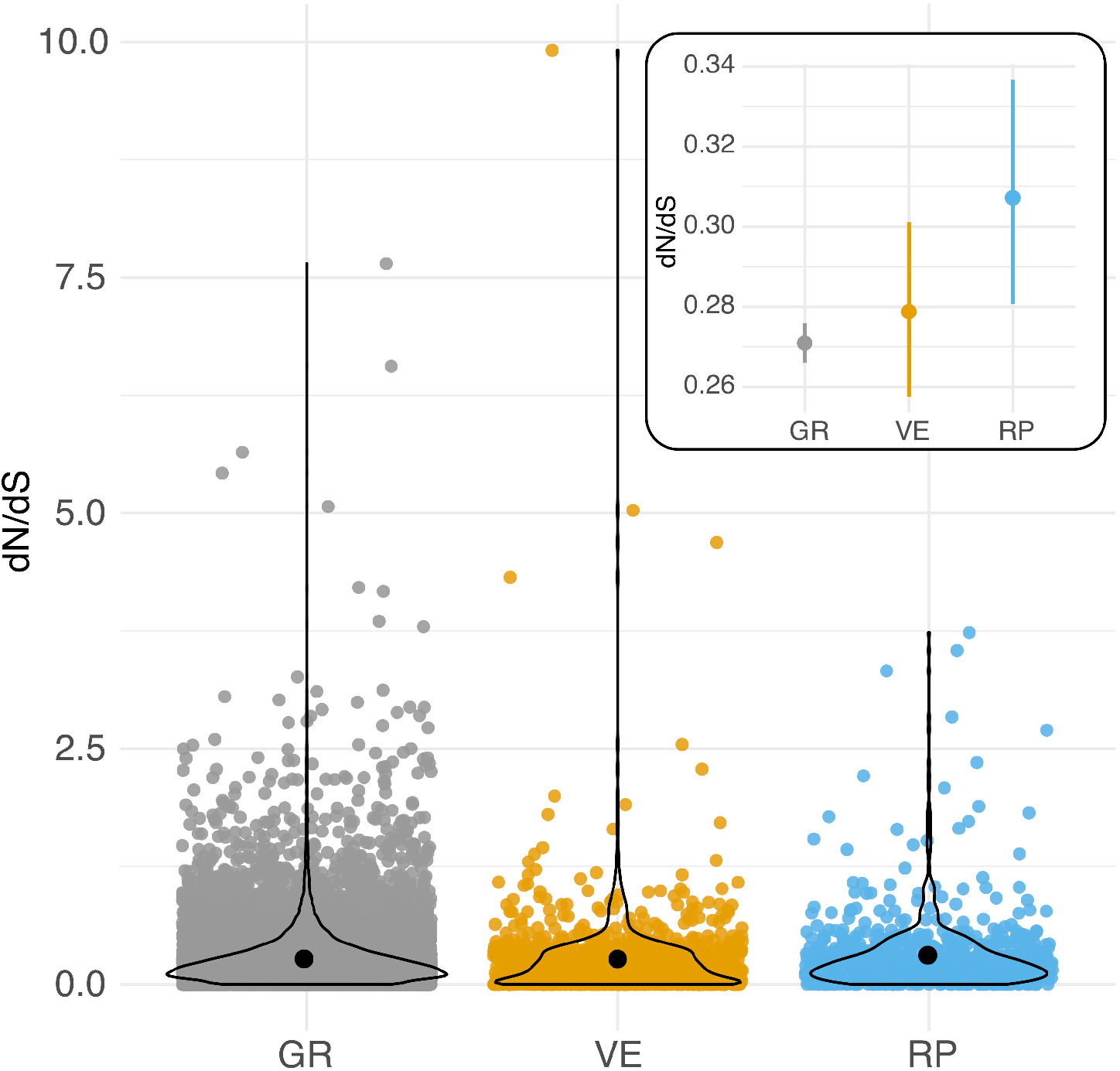
Per locus protein evolution (dN/dS) among loci expressed in reproductive (RP), vegetative (VG), and both reproductive and non-reproductive (general/GR) tissues in *Solanum* species. Black circles indicate means for each group. Inset: Estimated mean (circle) and 95% CI of dN/dS for each group of loci. Overlaid violin plots show the distribution of each group of data.

**Table 1.** Rates of protein evolution (dN/dS) in different broad classes of loci. dN/dS calculated for each locus using a one ratio test in PAML.

| Class | N loci | Per locus dN/dS |  |  | Tukey test<sup>1</sup> | Loci with dN/dS>1 |  |  |
| --- | --- | --- | --- | --- | --- | --- | --- | --- |
|  |  | mean | SE | Median |  | Number | Proportion<sup>2</sup> | Fisher's exact tests<sup>1</sup> |
| General | 16221 | 0.2709 | 0.0026 | 0.20 | a | 399 | 0.0246 | a |
| Vegetative | 923 | 0.2783 | 0.0111 | 0.19 | ab | 26 | 0.0282 | ab |
| Reproductive | 670 | 0.3071 | 0.0142 | 0.21 | b | 28 | 0.0418 | b |

1. see Table S4.
2. The proportion of genes with dN/dS>1 differed among classes of loci ( $X^2=7.9696$ , $p=0.0186$ ).

**Table 4:**
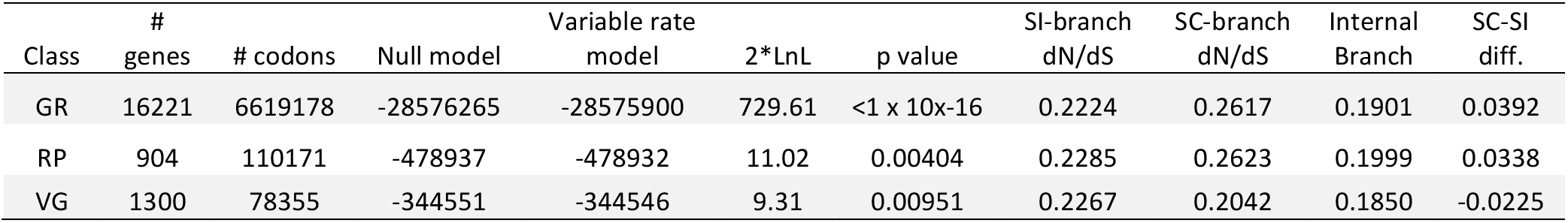
Estimates of dN/dS for SC and SI branches separately, for loci concatenated within each broad class of genes. Results are from 3 ratio model in PAML, with input species tree: ((Slyc #1, Spim #1), (Spen #2, Shab #2)).

### 3.2 Female- rather than male-specific genes evolve the most rapidly

Comparisons among tissue-specific genes indicated no evidence that pollen-specific loci evolve more rapidly than loci from female-derived tissues (style and ovule) (Figure 2; Table 2: Table S7). Instead, style- and ovule-specific loci had the highest mean dN/dS values of all tissue groups, and posthoc pairwise tests indicated that style mean rate was significantly greater than rates in root-specific and leaf-specific loci, and ovule mean rate was significantly greater than leaf mean rate (Figure 2; Table 2; Table S8). In comparison, pollen loci had a mean rate that was intermediate between these groups of tissue-specific genes, and statistically indistinguishable from each other tissue (Table S8). Note that many fewer loci were available for tissue-specific comparisons, especially leaf- and stem-exclusive loci (Table 2), and two tissues—vegetative meristem and seed—had no detected tissue-exclusive loci.

**Figure 2:**
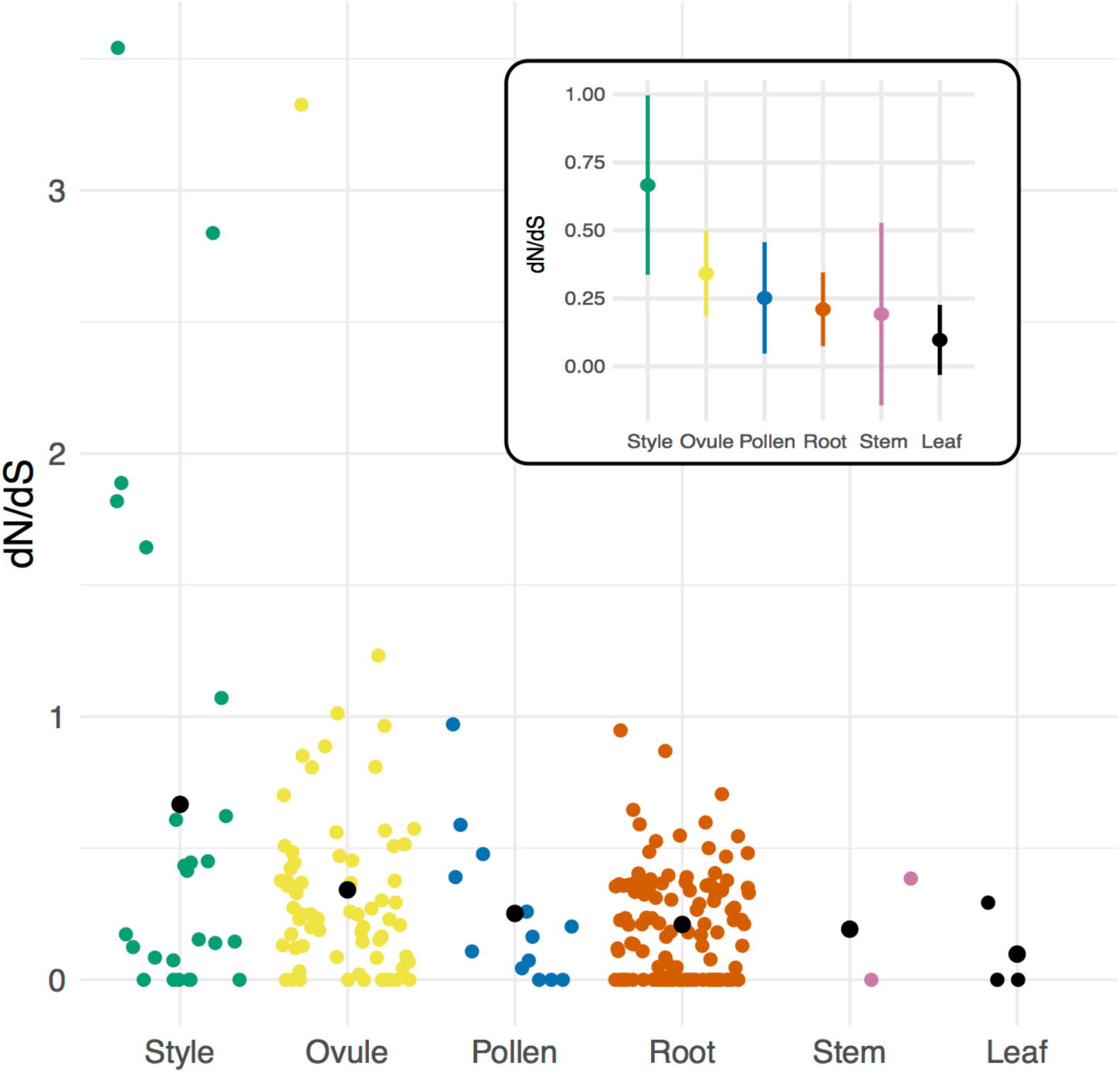
Per locus protein evolution (dN/dS) among groups of tissue-specific loci in *Solanum* species. Black circles indicate means for each group. Inset: Estimated mean (circle) and 95% CI of dN/dS for each group of loci.

**Table 2:**
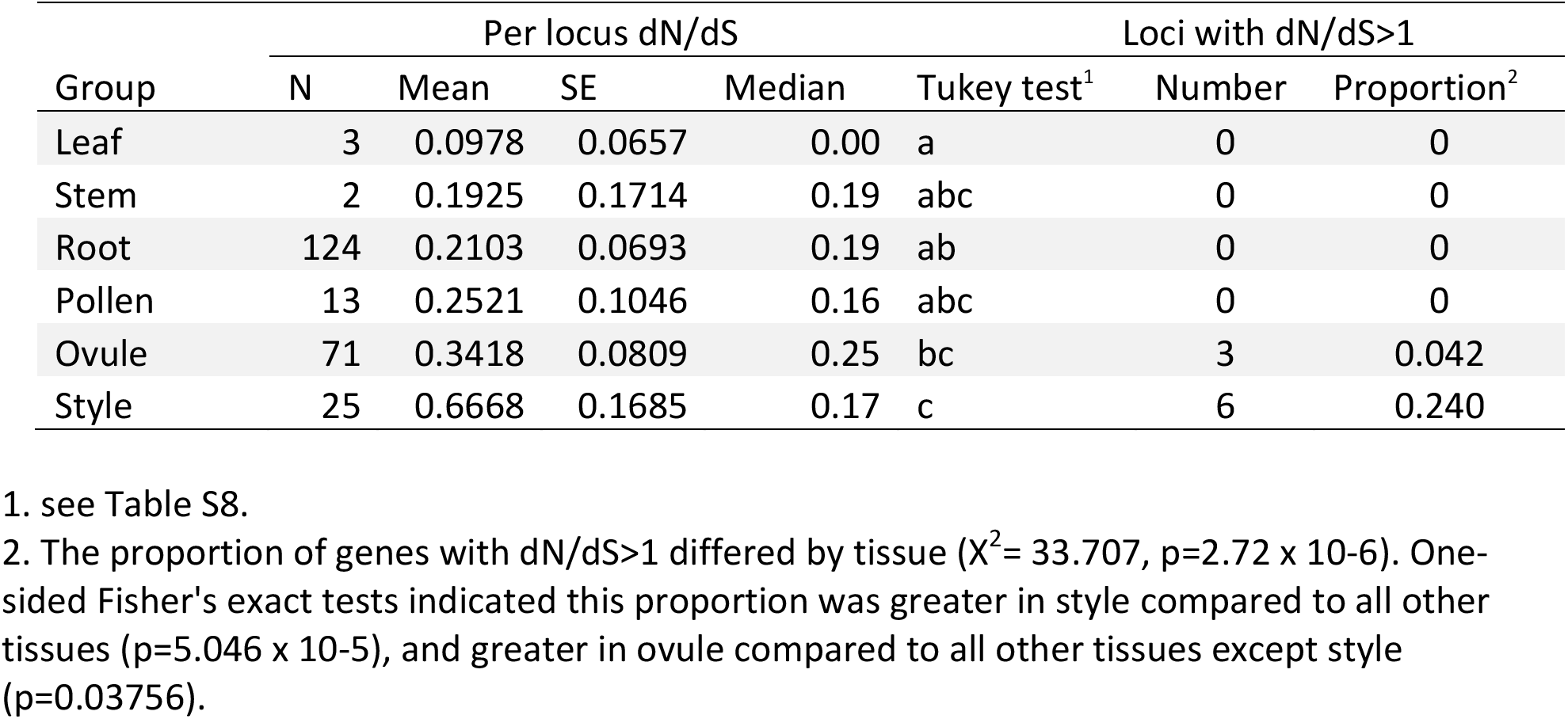
Rates of protein evolution (dN/dS) in tissue-specific loci.

Similar to comparisons of mean rates, the proportion of genes with dN/dS>1 differed by tissue (Table 2). Moreover, this proportion was significantly higher in style tissue—where 6 of 25 loci were inferred to be evolving adaptively—compared to all other tissues (Fisher’s exact test, p= 5.046 × 10^−5^; Table 2). For ovule-exclusive loci, the proportion adaptive evolving loci (3 of 71) was also greater than in all other tissues (excluding style-specific loci) (Fisher’s exact test, p= 0.0376) (Table 2).

### 3.3 Gametophytic-exclusive proteins do not evolve more rapidly than sporophytic-exclusive proteins, or proteins expressed in both tissue types

Although the mean rate of protein evolution in gametophytic genes was slightly higher than the rate of protein evolution in sporophytic-exclusive loci, this difference was not significant (t=-0.452, P=0.588) (Table 3; Table S9). Gametophytic-exclusive proteins also do not evolve faster than those expressed in at least one gametophytic and one sporophytic tissue (t=−1.498, P=0.134) (Table 3). Similar to mean rate comparisons, the proportion of gametophytic loci with dN/dS>1 (3 of 86, or 3.49%) did not differ from this proportion in sporophytic-exclusive genes (68 of 2033, or 3.34%), and was slightly but not statistically larger than this proportion in gametophytic+sporophytic genes (383 of 15778, or 2.43%) (Table 3).

**Table 3:**
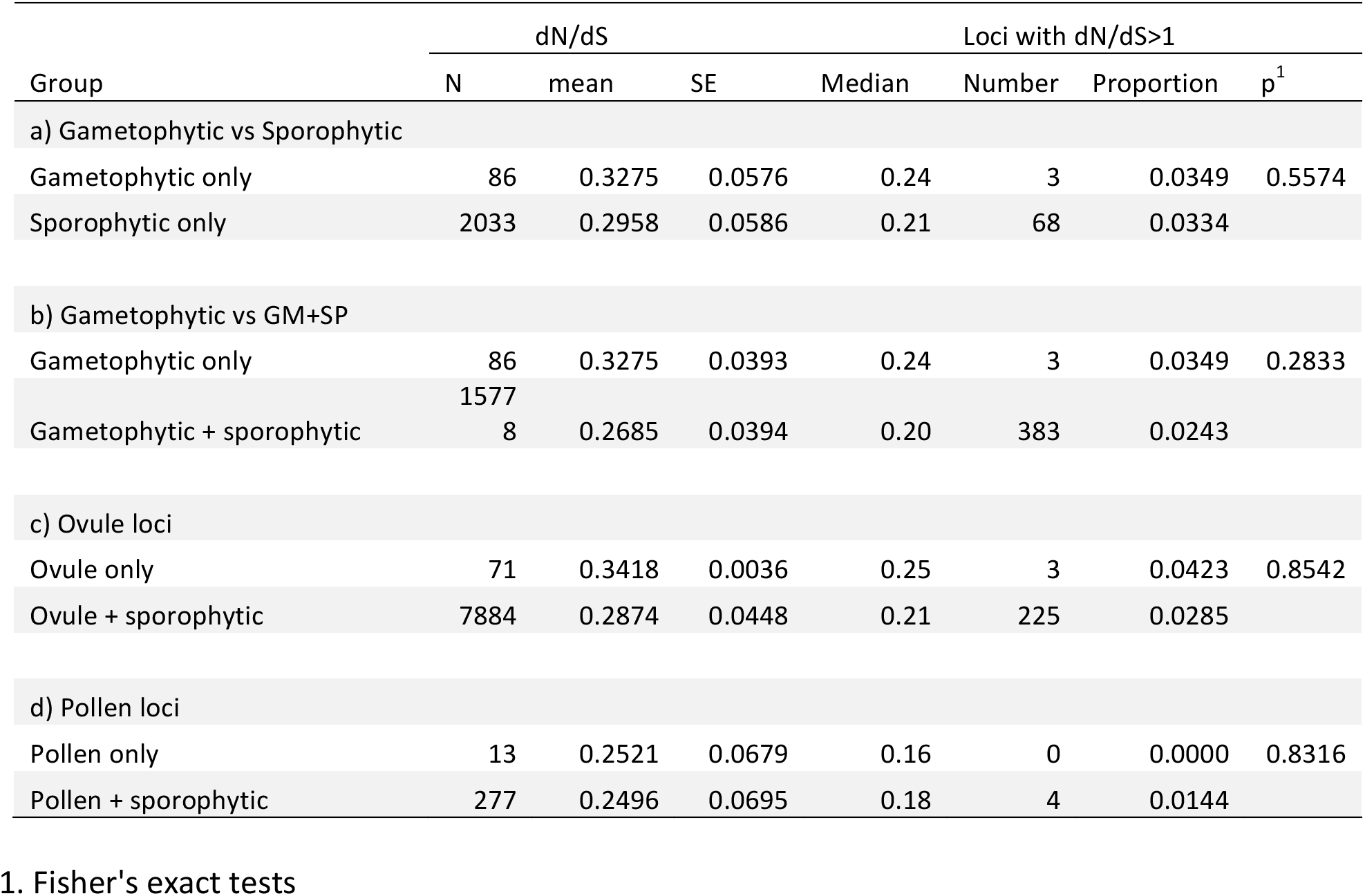
Rates of protein evolution (dN/dS) in gametophytic (GM) versus sporophytic (SP) or gametophytic+sporophytic (GM+SP) loci. Within each GLM, tissue groups did not differ in mean dN/dS (Table S9).

As observed in the general gametophytic comparisons, mean rate of protein evolution in ovule-exclusive loci was slightly higher than the rate of protein evolution in loci expressed in ovules and at least one sporophytic tissue (Table 3), but this difference was not significant (GLM t=1.212, p=0.225; Table S9). The proportion of ovule-exclusive loci with dN/dS>1 (3 of 71, or 4.23%) also did not differ from the proportion of ovule+sporophytic loci with dN/dS>1 (225 of 7884, or 2.85%) (Table 3). Neither the mean rate of protein evolution (GLM t=-0.037, p=0.971; Table S9) nor the proportion of loci with dN/dS>1 (Table 3) differed between pollen-exclusive loci and loci expressed in pollen and at least one sporophytic tissue.

### 3.4 Patterns of protein evolution show mating-system effects that are not specific to reproductive loci

Consistent with genome-wide effects of lower Ne in our self-compatible lineages, rates of protein evolution were inferred to be higher overall on self-compatible branches in both GR and RP classes of loci, although not for VG loci (Table 4); in the latter case, VG rates were numerically higher on SI branches. Results were similar for two-ratio models (Table S10). The observed difference in rates between SC and SI branch for RP loci (0.0338) did not fall outside the 95% CI of bootstrapped values (0.0007-0.0797) from the broader class of GR genes; instead 33% of bootstrapped samples had a dN/dS difference smaller than observed for RP loci.

### 3.5 Higher rates of protein evolution are consistently associated with greater tissue-specificity

Across loci classified according to the number of tissues in which they were expressed at >2 TPM (from 1 to 8 tissues), mean dN/dS was highest in loci expressed in a single tissue and consistently declined across loci expressed in increasingly more tissues (Figure 3; Table 5; Table S11, S12). Similarly, both mean expression level and tissue class (GR, RP, VG) affected the per locus estimates of protein evolution (dN/dS), as did their interaction (Table 6), indicating that the relationship between expression level and rates of protein evolution varies depending upon the broad tissue class. In particular, while the relationship between mean expression level and dN/dS was significantly negative for generally-expressed (GR) loci, the 95% CI for this slope overlapped zero for RP loci, indicating little evidence for a relationship between expression level and rate of protein evolution for the RP class of loci (Figure S3; Table S13). Classes of genes also differed in their overall mean gene expression (mean total TPM), which was highest in RP genes (505.2±23.7) compared to GR genes (398.7 ±4.2), and VG loci (43.9±4.5) (Table S14). Finally, mean gene expression level also differed between some tissue-specific loci, with style and ovule loci having the highest average gene expression (Table S15).

**Figure 3:**
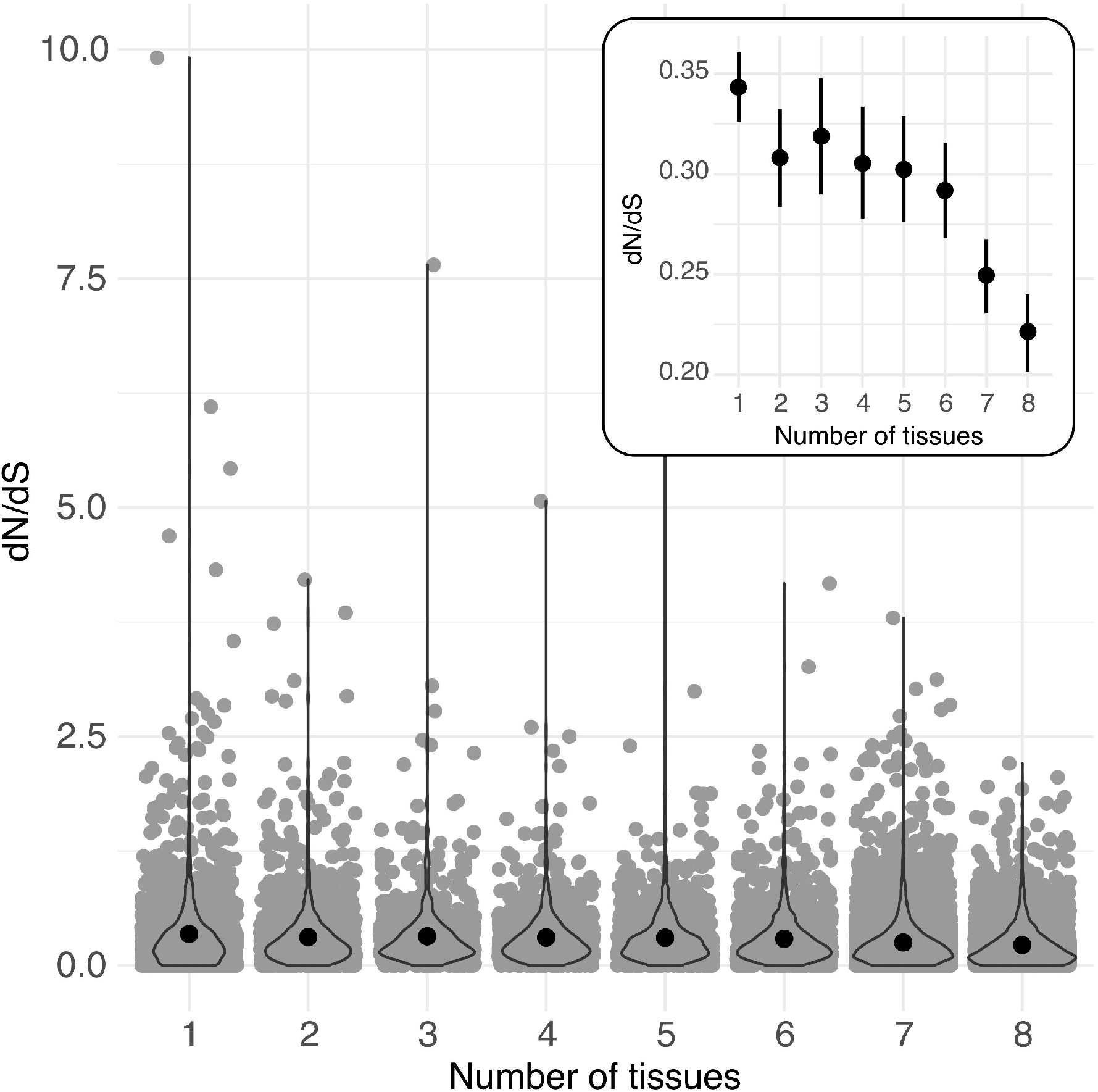
Per locus protein evolution (dN/dS) among groups of loci expressed (>2 TPM) in different numbers of tissues (from 1 to 8) in *Solanum* species. Black circles indicate means for each group. Inset: Estimated mean (circle) and 95% CI of dN/dS for each group of loci. Overlaid violin plots show the distribution of each group of data.

**Table 5:** Rates of protein evolution (dN/dS) in loci according to breadth of tissue expression.

| # Tissues<br>expressed in | N loci | Mean<br>sqrt(dN/dS) | SE | Median | Posthoc<br>Tukey test<sup>1</sup> |
| --- | --- | --- | --- | --- | --- |
| 1 | 1892 | 0.3429 | 0.0088 | 0.24 | d |
| 2 | 1528 | 0.3080 | 0.0125 | 0.22 | cd |
| 3 | 901 | 0.3183 | 0.0148 | 0.23 | cd |
| 4 | 930 | 0.3053 | 0.0142 | 0.23 | cd |
| 5 | 1083 | 0.3022 | 0.0135 | 0.24 | c |
| 6 | 1507 | 0.2919 | 0.0122 | 0.21 | c |
| 7 | 7206 | 0.2498 | 0.0094 | 0.19 | b |
| 8 | 3173 | 0.2213 | 0.0098 | 0.16 | a |

1. see Table S12

**Table 6:** Relationship between mean gene expression level and estimated dN/dS, across loci in each broad class (GR, RP, VG) of loci (Linear model)

|  | DF | SS | MS | F | Pr > F |
| --- | --- | --- | --- | --- | --- |
| Expression level | 1 | 20.53 | 20.5297 | 334.79 | 2.2 x 10<sup>-16</sup> |
| Class | 2 | 2.69 | 1.3467 | 21.96 | 2.98 x 10<sup>-10</sup> |
| Expression level x Class | 2 | 3.85 | 1.9239 | 31.37 | 2.50 x 10<sup>-14</sup> |
| Residuals | 17808 | 1092.02 | 0.0613 |  |  |

These findings are generally consistent with greater evolutionary constraint on loci with broader expression domains, across all our loci. In addition, for genes expressed in at least one reproductive and non-reproductive tissue (i.e., general loci), they indicate greater constraint on genes with higher magnitudes of gene expression.

### 3.6 Adaptively evolving reproductive proteins are not enriched for specific functional categories, but do include roles in sexual interactions

We found no evidence for enrichment of functional gene ontology categories among loci for which we found dN/dS>1, including among reproductive loci inferred to be adaptively evolving (data not shown). Nonetheless, among these reproductive loci, we did identify several genes with clear functional roles in sexual interactions (Table S16).

## 4 Discussion

Reproductive proteins appear to evolve rapidly in many animal groups. Here we evaluated evidence for elevated reproductive protein evolution in four closely-related flowering plant species, along with the potential influence of three factors on observed patterns: variation in mating system (and therefore the possible strength of sexual selection), in gametophytic gene expression (and therefore the strength of haploid selection), and in the breadth and/or magnitude of gene expression. We found evidence for elevated rates of reproductive protein evolution, globally across the genome and in some specific reproductive tissues. Moreover, these elevated rates appear to be more characteristic of female- rather than male-specific loci. Among the three broad factors that might influence these patterns, we found that elevated evolution was consistently associated with more narrow domains of gene expression, but not with expression in haploid versus diploid tissues; evolutionary rate differences between lineages with different mating systems (self-compatible versus self-incompatible) were more complex and not associated with specific shifts in reproductive protein evolution. Here we discuss these findings in light of previously observed patterns of reproductive protein evolution, and the factors proposed to drive these patterns.

### 4.1 Female reproductive proteins tend to evolve more rapidly

Our evidence for elevated rates of reproductive protein evolution was observed both as a modest but significant increase in mean dN/dS across all reproductive loci, and as a significantly larger proportion of loci showing evidence of adaptive divergence between species (dN/dS>1). Interestingly, however, this elevated evolution appears to be more clearly associated with proteins that have female- rather than male-specific functions. Indeed, female reproductive loci predominate among our most rapidly evolving loci, both generally and in analyses of tissue-specific loci. Of the tissue-specific loci found to have dN/dS>1, all are expressed in a female-specific tissue (either style or ovule) (Table S16).

This finding is intriguing. Some previous studies have detected accelerated evolution in proteins with specific female-associated functions, for example, egg proteins involved in sperm-egg interactions (e.g. Galindo et al. 2003). Nonetheless, most studies that include both male and female proteins have found mixed evidence for elevated female-biased or female-specific evolution (e.g. Wong 2010, Haerty et al. 2007; including in plants, e.g. Gossman et al. 2016), compared to much clearer accelerated evolution in male-specific or male-biased proteins (reviewed in Wong 2011, Wilburn and Swanson 2016). Some of this male-female difference could simply reflect a comparative lack of knowledge about female-associated proteins, leading them to be systematically underexamined in many comparisons. Indeed, most prior studies in animals (reviewed, for example, in Clark et al. 2006, Dapper and Wade 2020), as well as some in plants (Arunkumar et al. 2013, Harrison et al. 2019, and references therein), have focused largely or entirely on reproductive proteins with male-specific functions. Here, our loci were defined by tissue-specific patterns of gene expression, rather than from *a priori* expectations of function, which might have helped to ameliorate any such biases.

Our data also indicates a role for adaptive evolution in the observed acceleration of female protein divergence. Elevated rates of non-synonymous change (but values of dN/dS still less than 1) could result from relaxed purifying selection (see further below) or because positive selection has only acted on a subset of sites within each gene, or a combination of these effects. However, we also found a larger proportion of female (style and ovule) loci with dN/dS>1, suggesting a concentration of adaptive female protein evolution specifically involving post-pollination reproductive processes. One driver of this could be sexual interactions. In many animals, where female choice can be exerted both before and after mating (the latter as ‘cryptic female choice’) (Andersson and Simmons 2006), however in angiosperms male-female (pollen-pistil) interactions can only act after pollen arrives at a flower. Therefore, all evolutionary dynamics involving female choice or male-female interactions must act upon loci expressed in this relatively narrow post-mating window. Indeed, among our rapidly evolving (dN/dS>1) style loci, we detect at least one protein with putative functions in pollen interactions (a pistil-specific extension-like protein; Solyc02g078080), in addition to several biosynthetic enzymes (Table S16). (In contrast, 2/3 rapid ovule-specific genes have potential functions in stress response, possibly suggesting roles that are not directly related to sexual interactions—see below.) Moreover, in other angiosperms, ‘female’ loci that show rapid adaptive divergence are also mostly pistil-specific proteins involved in selection among different pollen genotypes (via roles in genetic self-incompatibility) (Clark et al. 2006), again suggesting the importance of post-pollination sexual dynamics in accelerated reproductive protein evolution.

Whether this potential adaptive explanation extends to explain the general elevation of female reproductive protein evolution (Figure 2) or reproductive proteins overall (Figure 1) is not yet clear. One prior study that compared protein evolution in male, female, and non-reproductive tissues in *Arabidopsis thaliana* and two relatives, also found that mean dN/dS was elevated in loci with expression biased towards female tissues (egg cell, central cell, and synergid cells: three haploid tissues within the female gametophyte) but not in pollen-biased proteins (Gossman et al. 2014). This pattern was attributed to relaxed selection on female-biased loci, partly because additional site-specific tests did not consistently detect evidence of positive selection in these loci (Gossman et al. 2014). Because our four species are closely related, we do not have sufficient power to assess site-specific evidence for adaptive evolution in loci whose estimated dN/dS is less than 1. Future analyses that pair divergence-based estimates with polymorphism data within species—that can provide alternative and often more powerful tests of positive and relaxed selection (e.g. Gossman et al. 2016, Arunkumar et al. 2013)—will be helpful in differentiating the relative influence of these different forms of selection on the broader patterns of elevated dN/dS detected here.

Regardless, if elevated female protein evolution is at least partly driven by post-pollination sexual interactions, it is interesting that we do not also observe elevated evolution in pollen-acting proteins. Of prior analyses in angiosperms, several have detected broadly elevated evolution in proteins that are pollen specific (e.g., Arunkumar et al. 2013, Gossman et al. 2016, Harrison et al. 2019). In principle, pollen protein evolution could be more constrained than style or ovule evolution due to factors like differences in the degree of tissue-specificity or in the operation of haploid selection, but these explanations are not generally consistent with our data (see further below). Alternatively, it’s possible that our analysis did not capture all the male proteins relevant to post-pollination sexual interactions. For instance, pollen loci whose expression requires direct interactions with the style and/or other pollen tubes would not be captured in our expression atlas (which was generated from RNA expression in mature ungerminated and *in vitro* germinated pollen; Table S1). There is evidence in other angiosperms that pollen transcriptomes can be actively modified during their transit through the style, resulting in changes in the suite of proteins being expressed (e.g., Qin et al. 2009, Mizukami et al. 2019). Accounting for and including these genes would require transcriptomes from ‘mixed’ post-pollination tissue, as well as specific approaches to differentiate pollen- from stylar-expression in these mixed samples (e.g. Pease et al. 2016b). Importantly, this limitation would also extend to any style- or ovule-specific proteins that are only elicited during pollination, so may not be a complete explanation for the general difference we detect between male and female loci here.

Other explanations for the under-accounting of pollen loci could include technical effects such as detection limits on gene expression in pollen, and/or exclusion of pollen-expressed loci that are especially short or rapidly evolving, due to filtering during transcript-mapping steps. In addition, by using dN/dS comparisons only, we would not have detected protein changes that involve duplicate, chimeric, or novel genes, and/or if different loci are the targets of selection in different lineages. If these latter factors do contribute to our failure to detect elevated pollen protein evolution, it suggests that different dynamics shape pollen protein identity and/or evolution in *Solanum* compared to other angiosperms that have shown evidence for elevated evolution using metrics like dN/dS (e.g. Arunkumar et al. 2013, Gossman et al. 2016, Harrison et al. 2019). Regardless, with the data that we currently have, we do not see evidence that male functions are subject to especially intense sexual selection, particularly in comparison to female reproductive proteins.

### 4.2 Rates of protein evolution are affected by tissue-specificity but not haploid expression

Our results can also suggest factors that shape the patterns of reproductive protein evolution that we do observe, at least among the three broad factors (gene expression level, haploid expression, and/or mating system variation) that we addressed here.

Of these, we find the strongest evidence that rates of protein evolution are influenced by the breadth and magnitude of gene expression. Many studies have shown higher rates of protein evolution in genes with more narrow expression domains and/or lower levels of gene expression (Meisel 2011, and references therein), a relationship that is thought to reflect lower constraint on tissue-specific proteins and stronger purifying selection on genes with high expression levels or broad expression domains. We similarly found that higher rates of protein evolution were associated with greater tissue-limitation and (in some cases) lower levels of gene expression in our species. However, for reproductive protein evolution specifically, we infer that tissue-limitation (a more narrow expression domain) is likely more important than magnitude of gene expression. For the latter, gene expression was not significantly associated with protein evolution for reproductive genes (Figure S3, Table S13), and reproductive loci also had much higher mean gene expression levels in comparison to generally-expressed genes and, especially, vegetative loci (Table S14).

High tissue-specificity has been proposed as an important contributor to elevated rates of reproductive protein evolution—and a better predictor of this than sex-biased gene expression *per se*—in both animals (Meisel 2011) and some plants (Veltsos 2019). For example, in *Arabidopsis thaliana* and close relatives, Harrison et al. (2019) found faster protein evolution in both pollen-specific and tissue-specific sporophytic genes compared to loci expressed in >1 tissue, although pollen-specific loci still retain the highest evolutionary rates in this and other analyses in these species (e.g. Szovenyi et al. 2013, Gossman et al. 2016). Arunkumar et al. (2013) also inferred a role for tissue-limitation or specificity in elevated dN/dS in *Capsella grandiflora* pollen proteins (although this elevation was attributed to different forms of selection—relaxed purifying selection in pollen germ cell genes versus greater purifying and positive selection in pollen tube loci—depending on the specific sub-tissues within pollen). These prior inferences, and our observations, suggest that higher tissue-specificity could facilitate broadly elevated rates of protein evolution. Nonetheless, tissue-specificity cannot provide a complete explanation of elevated reproductive protein evolution in our dataset, as tissue-specific non-reproductive proteins still evolve more slowly than tissue-specific (female) reproductive proteins (Table 2).

In comparison, we found no rate differences between loci expressed in haploid versus diploid tissues, and therefore little evidence for a strong effect of haploid selection in driving differences between reproductive and non-reproductive proteins. Haploid selection has two projected effects on haploid-expressed loci—increasing the efficacy of positive selection on advantageous alleles, and of purifying selection on deleterious alleles (Immler and Otto 2018, Mattila et al. 2018)—compared to diploid-expressed loci. We have limited ability to differentiate these two effects with our particular estimate of protein evolution, but our observations are not consistent with a strong effect of either. In terms of accelerating adaptive protein evolution, we found no significant elevation of evolutionary rates in loci limited to gametophytic-tissues in general, and in ovule-limited and pollen-limited loci specifically (Table 3). Similarly, we find little evidence for stronger or more widespread purifying selection on haploid-expressed loci, which would be observed as lower mean rates of protein evolution in gametophytic loci. Stronger purifying selection due to haploid expression has been proposed as one reason why sex-biased loci (generally, pollen expressed loci) do not appear to evolve faster than non-sex-biased loci in several dioecious species (Sanderson et al. 2019, Cossard et al. 2019; discussed in Muyle 2019, but see also Veltsos 2019) and, indeed, sometimes appear to evolve more slowly (e.g., Darolti et al. 2018). Our data do not suggest stronger purifying selection in either gametophytic-limited proteins in general, or ovule- and pollen-limited loci specifically, in our species. However, because we have relatively few pollen-specific loci (discussed above) we cannot exclude the possibility that intermediate rates of pollen protein evolution result from a tension between both positive and purifying selection predicted to act on these loci. Nonetheless, our present data indicates little evidence for a global effect of haploid gene expression on molecular evolutionary rates.

### 4.3 Mating system variation does not does not differentially influence reproductive protein evolution

Our results also did not support a strong consistent effect of mating system differences—and therefore predicted differences in the strength of sexual selection—on patterns of reproductive protein evolution. Interpreting these comparisons is complex because differences in genetic effective population size are expected to have global effects on rates of molecular evolution (Charlesworth and Wright 2001, and see Introduction) that are unrelated to sexual selection *per se*. The broad patterns of dN/dS we observed generally reflected these expected global effects of differences in Ne due to mating system variation: in the SC lineages, mean dN/dS was higher in both RP and GR genes, consistent with a general relaxation of selection in more selfing lineages (Charlesworth and Wright 2001) regardless of whether or not loci were associated with reproductive functions (Table 4). This genome-wide effect might have overwhelmed more subtle effects of mating system variation acting specifically on loci involved in male-male competition and/or female choice. In addition, the transition to self-compatibility is relatively recent (less than 1 MY) among our species, and self-compatible lineages still retain significant potential for outcrossing (Rick 1979, Vosters et al. 2014), which generally ranges from 0-40% (e.g. Rick et al. 1977) but has been estimated as high as 84% in some genotypes (Rick et al. 1978). These factors mean there may be only modest differences in the historical and current opportunity for sexual selection among our lineages, and therefore limited opportunity to observe systematic consequences of this transition as differential changes in evolutionary rates, especially among a sample of four species. (Statistically, this is compounded in our particular dataset by the two SC species branches being substantially shorter than those for our two SI species.) This contrasts with some other angiosperms, such as *A. thaliana*, where the transition to self-compatibility is about as old but the consequent effects on outcrossing rates has been much more severe (outcrossing rates are frequently <5%; Harrison et al. 2019, and references therein). Even among *Arabidopsis*, however, the effects of mating system on sexual locus evolution might be complex. For example, Gossman et al. (2014) found that dN/dS in pollen-acting genes was significantly lower in the outcrosser *A. lyrata* compared to selfing *A. thaliana*, even though pollen competition could be expected to accelerate adaptive evolution in these loci in the outbreeding species.

Interestingly, several previous analyses in animal systems have similarly found equivocal evidence for differential effects of mating system variation on the evolution of reproductive tract proteins (reviewed in Wong 2011). For example, in studies of primates, evolutionary rates in specific sperm and copulatory plug proteins are positively associated with common proxies for sexual selection, such as residual testis size and estimated number of mates (e.g. Dorus et al. 2004, Martin-Coello et al. 2009). However other studies find that phylogeny-wide signals of positive selection in reproductive tract proteins are not associated specifically with polyandrous (as opposed to monogamous) lineages (e.g., Ramm et al. 2008, Finn and Civetta 2010; and see Table 1 in Wong 2011). Limitations on phylogenetic models that can assess associations between molecular evolutionary rates and changes in phenotypic characters, and variation among lineages in the specific targets of sexual selection (Wong 2011), might explain relatively weak effects of mating system variation on protein evolution rates, despite the importance of sexual selection in driving faster reproductive protein evolution. Adding more species, and focusing on pairs of SI-SC sister species, might help in addressing some of these limitations in future analyses within *Solanum*.

Alternatively, the weak effect of mating system variation—both here and in prior studies— might be because the predominant cause of globally elevated reproductive protein evolution is not sexual selection. For example, pathogen resistance responses are also known to be associated with elevated rates of protein evolution—driven by antagonistic host-pathogen evolution—and might be concentrated among reproductive proteins if mating often leads to the exchange of pathogens (Clark et al. 2006, Wong 2011). In principle, the identity of loci inferred to be under positive selection could be helpful in differentiating among alternative selective agents responsible for driving this adaptive divergence. Of the genes in our dataset that show elevated evolution, we cannot yet draw strong conclusions about whether they better reflect roles related to sexual interactions versus responses to non-sexual environmental factors; many of them have unknown functions or general functions that do not *a priori* provide specific support for either inference (Table S16). We do detect at least one style-specific protein with a clear functional connection to pollen-pistil interactions. However, our rapidly-evolving ovule-specific genes are implicated in stress responses, which might indicate a greater role for natural rather than sexual selection in shaping adaptive responses in this class of loci.

### 4.4 Consequences of rapid protein evolution for the evolution of reproductive differences and speciation

Finally, among the motivations for examining patterns of reproductive protein evolution is to understand their possible role in lineage diversification. If reproductive proteins do indeed evolve more rapidly than other classes of protein, then perhaps they play an outsized role in the emergence of functional differences and reproductive isolating barriers between lineages. For example, evidence suggests post-mating prezygotic traits frequently evolve rapidly in animals (e.g. Galindo et al. 2003, Dorus et al. 2004, Plakke et al. 2019) and, when divergent between species, could contribute to reproductive isolation (Howard 1999, Snook et al. 2009, Palumbi 2009, Castillo and Moyle 2014). Similarly, divergence in post-pollination prezygotic reproductive molecules could be important contributors to species barrier formation among angiosperms (Moyle et al. 2014). Here we did detect evidence for a global elevation of protein evolution in tissues that mediate these post-pollination prezygotic stages, especially of female-specific loci. Our observations are consistent with a role for tissue specificity in facilitating these elevated rates, but not for haploid selection or, as yet, for differences in selective dynamics that are associated with recent shifts in mating system. The evolutionary factors that might drive this more rapid evolution therefore remain to be clarified. Regardless, our results indicate that rapid reproductive divergence is especially characteristic of proteins with female-specific functions among our species. These specific functions can be critical for maintaining coordinated sexual signals and ensuring fertilization post-pollination (Bernasconi et al. 2001, Swanson et al. 2004, Moyle et al. 2014). Therefore their rapid divergence might be particularly influential in the initial stages of reproductive isolation and, thereby, in speciation, in this highly diverse plant genus.

## 5 Author contributions

LCM, MW, and MJSG designed the study. MW and MJSG performed the analyses. LCM wrote the manuscript with assistance from MW and MJSG.

## 7 Acknowledgments

Thanks to CJ Jewell for collecting ovule tissue, Christie Bergeron for RNA extractions, the Indiana University Center for Genomics and Bioinformatics for assistance with library construction and sequencing, and three anonymous reviewers for comments that helped improve the study.

## Supplementary Figures

**Supplementary Figure 1:**
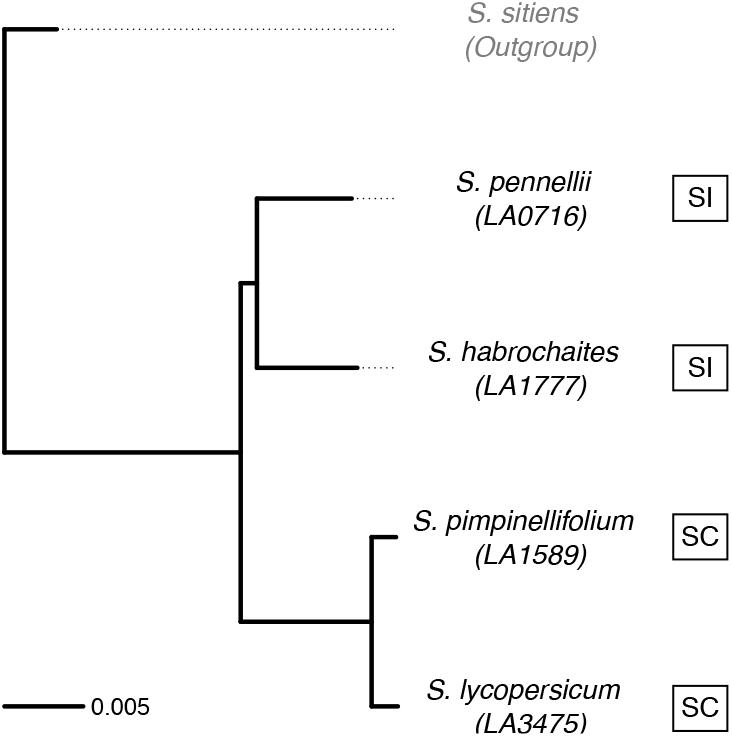
Maximum likelihood tree of focal species relationships relative to an outgroup species *Solanum sitiens*. Branch lengths (in average substitutions per site) were obtained from Pease et al. (2016a). Original tree was inferred with RAxML v.8 (Stamatakis, 2014) from a whole-transcriptome concatenated alignment.

**Supplementary Figure 2:**
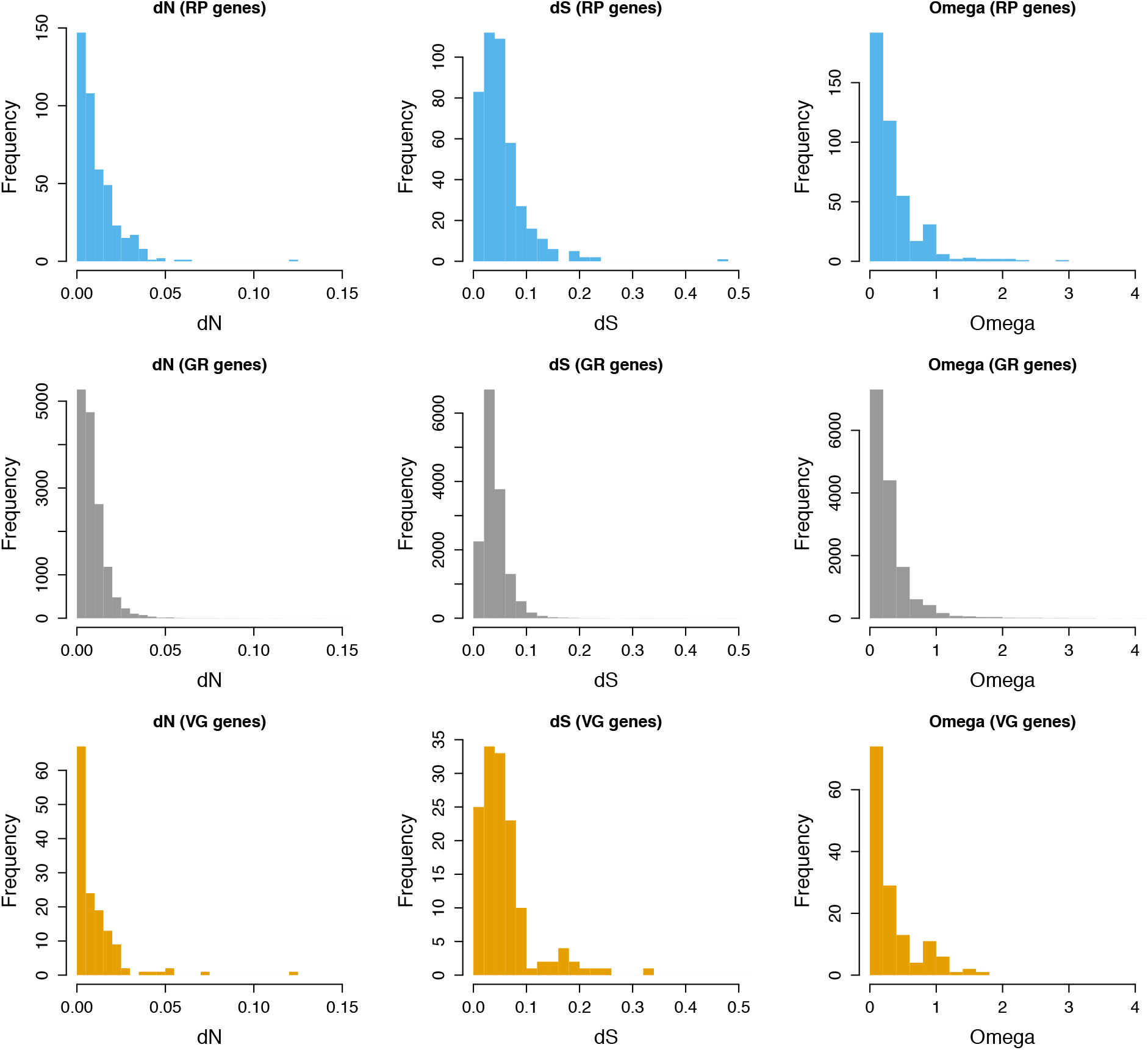
Distributions of inferred per-locus dN (left), dS (middle), and dN/dS (omega) (right) from each broad class of loci: RP (upper panels), GR (middle panels), and VG (lower panels). Supporting analyses are in Table S6.

**Supplementary Figure 3:**
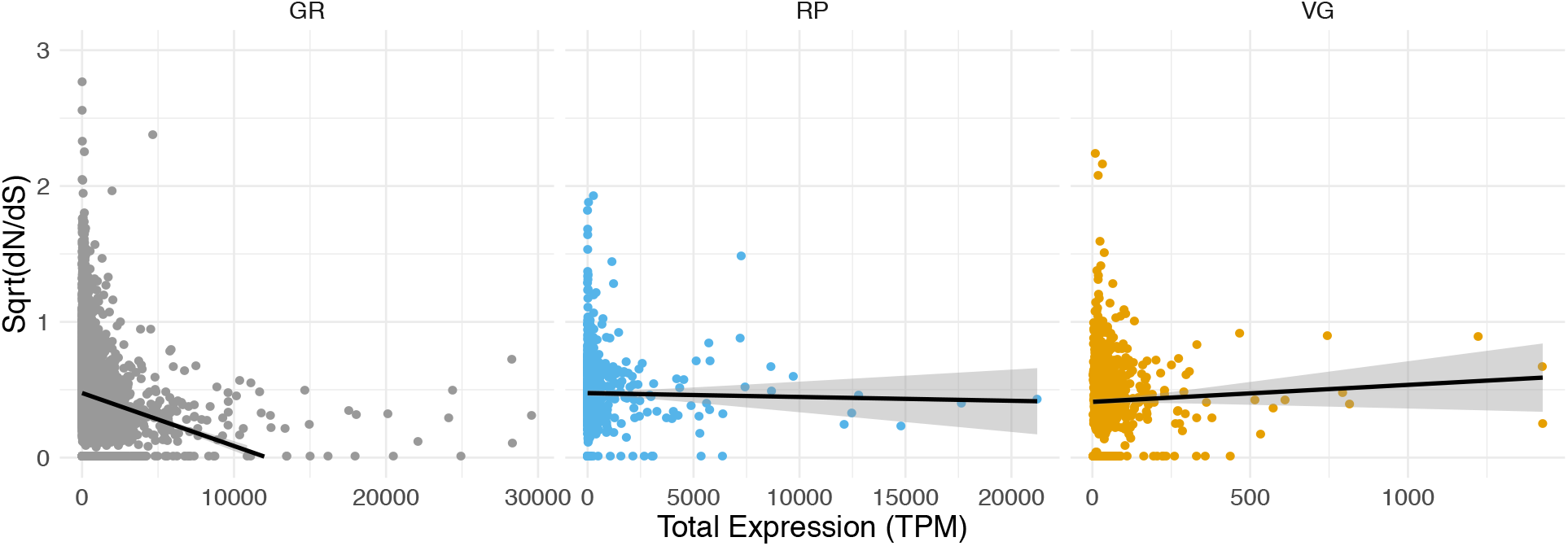
Relationship between per locus mean level of gene expression (TPM) and per locus protein evolution (dN/dS) in loci expressed in (left) general/GR, (middle) reproductive/RP, and (right) vegetative/VG tissues in Solanum species. Each point indicates an individual locus. Regression lines shown in black; 95% CI shown in grey. Supporting analyses are in Table S13.

